# Retreat Site Selection in *Vaejovis carolinianus* Populations of Tennessee’s Upper Cumberland Region

**DOI:** 10.1101/058354

**Authors:** Bob A Baggett

## Abstract

My study examined Tennessee’s only native scorpion species, *Vaejovis carolinianus*. Little is known about its ecology, so the objectives of my study were to determine if: (1) *V. carolinianus* selected cover objects based on surface area; (2) *V. carolinianus* preferred moister soils under the cover object; and (3) length of time in captivity altered the preferences. Scorpions were captured from two different locations: (1) a roadcut parallel to State Highway 96 near Edgar Evins State Park; (2) France Mountain in Overton County. In laboratory trials, scorpions were allowed to choose among three retreat sites and three soil moisture levels. Transects were established at both field locations to count and measure rocks that may serve as retreat sites. Surface area trials indicated that *V. carolinianus* selected large objects as retreat sites most often, but overall retreat site selection did not differ from that expected based on random choice weighted by cover object size. Soil moisture trial results varied, with no statistical significance in the results from the Highway 96 and 2013 France Mountain populations. The 2014 France Mountain population did show statistically significant differences. The surface area trials did not exhibit a time in captivity effect, but the soil moisture trials indicated a time in captivity effect between the 2013 and 2014 France Mountain populations. Based on test results, it appears *V. carolinianus* selected larger rocks as cover sites, perhaps due to chance or perhaps to escape sunlight and heat, for predator avoidance, or for higher soil moisture levels.

## INTRODUCTION

The study of how and why organisms select particular habitats has long been central to ecology (Huey, 1991). Many factors contribute to the habitat selection process, including environmental factors (temperature, precipitation, soil type, etc.) occurring at retreat sites; availability of shelters, food, nest sites, and mates; abundance of conspecifics and competitors of other species; and risks of predation, parasitism, and diseases (Hodara and Busch, 2010). Retreat sites are locations selected by organisms for the degree of safety they offer from predators, accessibility of food, and/or appropriate thermal or humidity regime (Langkilde and Shine, 2004). Many animals spend long periods of time within retreat sites, and individuals are often philopatric to such sites (Croak et al., 2008). Selection of thermally suitable retreat sites may be particularly important for ectotherms, because behavioral and physiological processes of these animals depend strongly on temperature (Downes and Shine, 1998). Availability of suitable retreat sites may have strong impacts on individual fitness as well as population viability. Accordingly, animals select retreat sites non-randomly and often use multiple criteria to prioritize suitable sites (Croak et al., 2008).

Relatively little is known about which factors influence retreat site choice in scorpions. *Vaejovis electrum* and *V. cashi* select large, thick rocks, probably due to lower temperature difference and lower daily thermal variance than smaller, thinner rocks (Becker, 2013). Abushama (1964) found negative reactions to light and hygro-positive behavior lead the scorpion *Leiurus quinquestriatus* to holes and crevices under stones and debris in open desert where conditions were suitable for survival during the day time.

Two species of scorpions are found in Tennessee, but only one, *Vaejovis carolinianus*, is native to the state. *Vaejovis carolinianus* has many common names, but the two commonly used are Plain Eastern Stripeless Scorpion and Southern Devil Scorpion. It is reddish to dusty-brown, up to 6.5 cm in total length, and can live for seven to eight years (Benton 1973). *Vaejovis carolinianus* inhabits the southeastern portion of the USA where it commonly resides near the edge of moist forest habitats (Croak et al., 2012). Diurnally, they are usually located under rocks, leaf litter, or bark of dead trees. This species also occurs around human dwellings if sufficient cover is available. Unlike other species in its family, *V. carolinianus* is a weak burrower, and it uses retreat sites with natural spaces or crevices (Benton, 1973). *Vaejovis carolinianus* is most active above 25°C and sluggish in colder temperatures (Elston, 2006). The species is viviparous with a gestation period of one year.

Male and female *V. carolinianus* have different habitat preferences (Benton, 1973; Dobbe, 1999). Adult females typically occur under and among rocks, as well as at the base of dead standing trees. Males and immature scorpions occur most commonly in leaf litter, as well as under the bark of dead trees (preferably pine). Mating occurs in late August; otherwise, adults do not usually interact (Polis, 1990; Dobbe, 1999).

The objectives of my study were to test several hypotheses on habitat preferences of female *V. carolinianus.* (1) Does the size (area) of a cover object influence its choice as a retreat site? (2) Do these scorpions have a preferred level of soil moisture? (3) Does length of time in captivity affect preferences for cover object size or soil moisture?

## MATERIALS AND METHODS

*Scorpion Collection and Care:* I captured a total of 110 *Vaejovis carolinianus* (99 female, 11 male) from two locations in Tennessee’s Upper Cumberland region during June through September 2013. My first location was a roadcut (36.120077° N, 85.811714° W, altitude 217.06 m) along Tennessee State Highway 96 (Hwy 96) leading to Edgar Evins State Park, approximately 2 km south of the intersection with Interstate 40. My second location included logging roads and electric line accesses off of Pleasant Shade Lane on France Mountain (36.240359° N, 85.237161° W, altitude 393.32 m) in Overton County, TN. The logging road entrance was 4.67 km from the intersection of Pleasant Shade Lane and Dry Hollow Road, and the electric line access point was 4.99 km from the same intersection. These locations were impacted by timber-cutting and rock collection. I captured scorpions during the day by turning over rocks and logs along edges of moist forests and in road cuts. Upon capture, scorpions were placed in 60 mL plastic bottles or 50 mL centrifuge tubes for transport back to Tennessee Technological University (TTU).

I captured thirty additional scorpions from France Mountain locations during May and June 2014. Instead of capturing *en masse*, six scorpions were captured initially, followed by an additional three individuals every two days until a total sample size of 30 scorpions was reached. I used this technique so all scorpions started lab trials within three days of capture.

I individually weighed (in mg) and measured (total body length minus telson and aculeus, in cm) each scorpion, and determined its sex by examining the pectines. These comb-like sensory organs are located ventrally and larger on males as well as contain more teeth. I recorded total length, weight, sex, and a unique identifier number on an Excel spreadsheet. Scorpions were placed individually in 17 cm × 11 cm × 5.8 cm plastic containers containing a 1 cm deep layer of sand substrate with a square (~4 cm^2^) of moistened paper towel for water and cover. I fed each scorpion one banded cricket, *Gryllodes sigallatus* of the same size as the scorpion or smaller, every seven to ten days while in captivity. Females with juveniles on their backs were not fed crickets until juveniles were independent (i.e., no longer on mother’s back). After offspring dispersal, juveniles were placed in separate containers and the female was immediately fed a small cricket. I checked paper towels every other day to maintain moisture and to check for mold. I replaced paper towels showing any evidence of mold.

Prior to and between trials, scorpions were stored in separate rooms during daylight and nighttime hours to approximate natural light:dark cycles (14:10 summer, 10:14 winter). Mean ambient temperature of both rooms was 24° C - 27° C. All females were assumed to be gravid at the time of capture (Benton, 1973), and 87 of 99 females gave birth between August and October 2013. To avoid potential biases in behavior associated with differences in timing of gestation, experimental trials began only after females had given birth.

*Field Procedures:* During July 2014, I established transects at both study sites to determine abundance of rocks of different size classes and preferred rock sizes used by scorpions as retreat sites. At the Hwy 96 site, I randomly established one 10 m transect parallel to the south side of the road, approximately equidistant between the edge of the road and the bluff. At France Mountain sites, I established three transects: a 10 m transect along an electric line access on which a fire pit was located, and two 10 m transects along a logging/rock-collecting access road. The fire pit was 0.40 km north of Pleasant Shade Lane and 14 m from the forest edge. The first transect on the logging/rock-collecting access road ran south to north, 25 m from and parallel to Pleasant Shade Lane, while the second transect started from the same point but ran in a west to east direction down the mountainside. I measured (length and width in cm) all rocks ≥ 85 cm^2^ touching or falling within a 1-m wide strip centered on transects. I counted rocks with a surface area ≤ 85 cm^2^ but did not measure them, as I determined these were too small to be an adequate retreat site. Additionally, I identified rocks as being embedded or not because *V. carolinianus*, a weak burrower, would be unable to crawl under embedded rocks. I flipped over all non-embedded rocks within transects and recorded whether they were being utilized as a retreat site or not. Due to lack of scorpions at the Hwy 96 transect, more than100 randomly selected rocks outside the transect were flipped over, including rocks from across the size range found in the transect. A subset of these rocks (38 in total) was measured for surface area (except for those less than 85 cm^2^), as were all rocks found with a scorpion beneath them.

*Laboratory Procedures*—I started experimental trials during the first week of October 2013. All trials were conducted in a Percival Intellus Model I36LLC8 Environmental Controller in order to control ambient temperature and light. Since the environmental controller did not have an outside water source, I maintained humidity by placing a two-gallon stock pot full of water on the controller’s bottom shelf. Each trial was conducted at 27.8° C and 34% relative humidity. I selected these values based on average summer daytime temperature and humidity levels for middle Tennessee. I placed scorpions in the environmental chamber set to a light:dark cycle of 14 hours light:10 hours dark, since this resembled the normal summer daylight pattern in Tennessee. I constructed test trays 67.3 cm long × 73.7 cm wide × 15.2 cm high from 0.9 cm thick plywood. I lined the tray walls with sheet metal to prevent scorpions from escaping and filled each tray with a 3.8 cm thick layer of soil. Substrate was changed and tiles were cleaned with dishwashing liquid then wiped with ethanol after each trial to remove potential chemical cues confounding results of subsequent trials. I placed scorpions, still in their individual containers, in the environmental chamber 72 hours prior to the start of each experimental trial to acclimate to test conditions.

The first set of trials examined cover object surface area preference. These trials used square ceramic tiles of three sizes: 232 cm^2^ (Small), 522 cm^2^ (Medium), and 929 cm^2^ (Large). Three tiles (one of each size) were arranged in a triangular pattern in each tray. I placed a scorpion in the center of the tray at the beginning of the light period. The scorpion remained undisturbed for one light:dark cycle. At the end of the cycle, I lifted tiles, located the scorpion, and recorded selected tile size. To assess repeatability of preferences, I repeated trials twice more for each scorpion, giving each three consecutive days of preference data. After each trial, I removed the scorpion from the tray, reoriented tiles by rotating the pattern in a clockwise direction, and placed the scorpion back in the center of the tray. At the end of three trials for an individual scorpion, I replaced tiles and substrate before another scorpion was placed in the tray.

The second set of trials examined scorpions’ preference for moisture content of soil. I divided the tray into three sections using 3.81 cm high partitions constructed from 0.9 cm thick plywood, equaling depth of the substrate. I filled each section with substrate of different moisture content (by percent moisture) and placed a single 232 cm^2^ tile at one end of each section.

Moisture content of soil was determined by first weighing a 1000 ml glass beaker on a digital scale, zeroing the scale, and then adding 500 g of the dried soil mixture. Using the formula below, enough

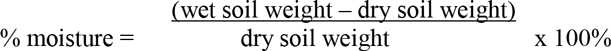

I repeated this process until each section of all three trays was filled to a depth of 3.81 cm with soil at the assigned moisture range. Twelve hours into each daily trial, I took a 50 ml soil sample from a randomly selected area in each moisture section to verify soil remained within assigned moisture ranges. I used the formula above to compare weight of a 50 ml sample to weight of a 50 ml sample of dry soil mixture. If the sample was outside the assigned range, I sprayed the entire section with tap water and took another randomly selected sample.

Moisture content was divided into three ranges: 0 – 15% (Dry), 16 – 30% (Damp), and 31 – 45% (Wet). I used ranges in this order inside each tray throughout the entire trial period. I used moisture ranges instead of individual values because the environmental controller did not have an external water source, so humidity level inside could not be automatically maintained.

Substrate used in test trays was composed of a mixture (50:50) of potting soil and top soil. This mixture was used to minimize weight of the trays which, in turn, minimized wear and tear. In order to sterilize the substrate, I spread it onto cookie sheets and dried for 12 hours in a conventional oven set at 110° C and allowed to cool for 30 minutes before sealing it in plastic Ziploc^®^ bags for transport back to TTU for use. I sterilized the substrate prior to beginning trials as well as after each scorpion’s three consecutive trial period.

At the beginning of a light:dark cycle, a scorpion was placed at a randomly selected section (selected by die roll) in the tray at the end opposite the tiles. The 0-15% soil moisture section was selected with a die roll of 1 or 4; 16-30% was selected with a die roll of 2 or 5; and 31-45% was selected with a die roll of 3 or 6. At the end of the 24 h cycle, I lifted tiles, located the scorpion and recorded its soil moisture selection. Additionally, I checked soil moisture content to ensure it was still within the correct percent moisture range. I removed the scorpion from the tray, reoriented tiles, and made and necessary moisture adjustments. Tile reorientation was accomplished by shifting each tile one section to the right, and the tile on the right end was moved to the section on the left end. The scorpion was placed back into a randomly selected section of the tray for two more consecutive 24 h trials. After three consecutive trials, I replaced tiles and substrate before another scorpion was placed in the tray.

Experimental trials with scorpions captured during the 2013 field season were completed in early May 2014. After repairing test trays, I started trials with scorpions captured during the 2014 field season (scorpion capture and experimental trials occurred concurrently). I used the same experimental procedures with the 2014 scorpions that I used with the 2013 scorpions, in the same order. The purpose of these procedures was to determine if scorpions collected in 2013 became acclimated to their state of captivity.

*Data Analyses:* Data collected in size preference, moisture content, and aggregation trials were compared both within and among the three populations (Hwy 96 roadcut, France Mountain, and France Mountain 2014). I analyzed my data using Statistical Programming for the Social Sciences (SPSS) version 21.0 (SPSS, 2012) with significance set at α = 0.05 (two-tailed).

I ran two chi-square goodness-of-fit tests (*G*-tests) to compare choice of tile sizes within the three populations. The first *G*-test compared observed values (choices) to expected values under the assumption each tile had an equal chance of being selected (expected values equal). The second *G*-test compared observed values to expected values that were based on tile surface area, such that expected probability of selecting a tile equaled the surface area of that tile divided by the total surface area of all three tiles. For these analyses, observed *N* was obtained by treating each trial day as a separate event; thus, each scorpion contributed a maximum of three choices to this total. Each scorpion’s daily tile choice was also assigned a score (Small = 1, Medium = 2, Large = 3), and these scores were summed for a total selection score.

Paired samples *t*-tests were used to compare observed total selection scores to expected total selection scores. Expected total selection scores were calculated by assuming each scorpion would select a different-sized tile each day of the three-day trial period (expected total selection score = 6). Using observed total selection score as the dependent variable, linear regression was used to determine if scorpion length affected tile size selection. I used a chi-square contingency table with surface area choice as the dependent variable to determine if there were any differences in surface area choice among the three populations.

I classified rock data collected from transects into one of five categories: (1) Very Small (VS) (< 85 cm^2^); (2) Small (S) (85 – 315 cm^2^); (4) Medium (M) (315 - 545 cm^2^); (5) Large (L) (545 - 775 cm^2^); and Very Large (VL) (> 775 cm^2^). Categories (with the exception of Very Small) were determined by dividing area measurements of rocks greater than 85 cm^2^ into 4 categories with equal area ranges. Initially, I combined rock measurements across all four transects, and used a Mann-Whitney *U* test to compare sizes of occupied and unoccupied rocks. I used a *G*-test to compare number of occupied rocks in each size class to total number of rocks available in each size class. Expected *N* values were adjusted based on rock size by entering the three tile surface area sizes into the “value” block of non-parametric chi-square analysis in SPSS. Finally, I compared rock size distributions among the three transect locations using a Kruskal-Wallis non-parametric ANOVA.

I ran *G*-tests to compare choice of soil moisture content within the three populations. Observed *N* was obtained by treating each trial day as a separate event; thus, each scorpion contributed a maximum of three choices to this total. Each scorpion’s daily moisture choice was assigned a score (Dry =1, Damp = 2, Wet = 3), and these daily scores were summed for each scorpion’s total moisture selection score. Using paired samples *t*-tests, I compared observed total moisture selection scores to expected total moisture selection scores. Expected total moisture selection scores were determined by assuming each scorpion would select a different moisture level each day of the three-day trial period (expected total moisture selection score = 6). Using observed total moisture selection score as the dependent variable, linear regression was used to determine if scorpion length affected soil moisture selection. I used a chi-square contingency table to determine if there were any differences in soil moisture choice among the three populations.

## RESULTS

*Choice of Surface Area of Retreat Sites:* High mortality prior to the beginning of trials resulted in only female scorpions being tested because all males had died (along with many females). Population sizes were reduced to: Hwy 96 roadcut (*n* = 24), 2013 France Mountain (*n* = 46), and 2014 France Mountain (*n* = 30). Two Hwy 96 roadcut scorpions were found elsewhere in the tray (not under a tile) on a single trial day; therefore, I excluded their trials from the analyses.

*Vaejovis carolinianus* from the Hwy 96 roadcut population displayed significant preference for large tiles when each tile was treated as being equally likely to be selected (*χ*^2^ = 10.400, *P* = 0.006, df = 2, expected *N* = 23.3). When expected values were scaled by tile surface area, no significant preference was found (*χ*^2^ = 1.360, *P* = 0.507, df = 2; expected *N* values were Small = 9.7, Medium = 21.7, and Large = 38.6). Although no significance was found with this second analysis, residual values indicated that large tiles were selected slightly less than expected (residual = −4.6) while small and medium tiles were selected slightly more than expected (residual = 2.3 for both) (Table 1; Figure 1). A *t*-test revealed a significant difference (*t* = 2.429, df = 24, *P* = 0.023) between observed tile size selection scores and expected tile size selection scores with observed occurring more than expected. Linear regression showed no significant relationship (*r*^2^ = 0.085, F_1,22_ = 2.051, *P* = 0.166) between observed tile size selection scores and scorpion length.

**Table 1:** Observed and expected *N* values with residual values (observed – expected *N*) from chi-square tests on tile size selection with relative surface area included in calculations of expected values.

| Population | Tile Size | Observed $N$ | Expected $N$ | Residual |
| --- | --- | --- | --- | --- |
| Hwy 96 Roadcut | Small | 12 | 9.7 | 2.3 |
|  | Medium | 24 | 21.7 | 2.3 |
|  | Large | 34 | 38.6 | -4.6 |
| 2013 France Mtn | Small | 30 | 18.9 | 11.1 |
|  | Medium | 44 | 42.5 | 1.5 |
|  | Large | 63 | 75.6 | -12.6 |
| 2014 France Mtn | Small | 11 | 12.3 | -1.3 |
|  | Medium | 25 | 27.6 | -2.6 |
|  | Large | 53 | 49.1 | 3.9 |

**Figure 1:**
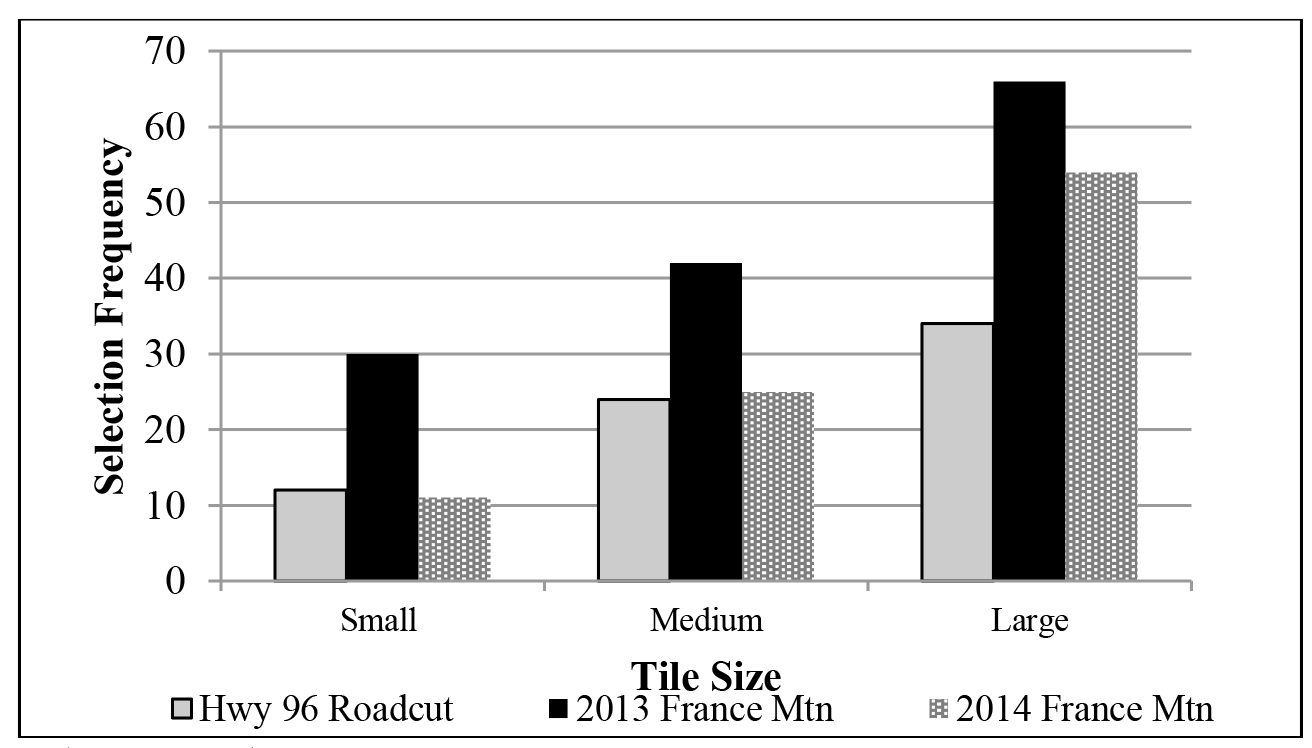
Tile surface area selection frequency of scorpion populations captured from two separate locations over two years

For the 2013 France Mountain population, *V. carolinianus* displayed significant preference for larger tiles when each tile was equally likely to be selected (*χ*^2^ = 12.015, *P* = 0.002, df = 2, expected *N* = 45.7). When surface areas were included in analysis, significant preference was found among the three tile sizes (*χ*^2^ = 8.671, *P* = 0.013, df = 2). Expected *N* values for small, medium, and large tiles were 18.9, 42.5, and 75.6 respectively. When examining residual values from the second *G*-test, scorpions selected small tiles more than expected (residual = 11.1), medium tiles slightly more than expected (residual = 1.5), and large tiles less than expected (residual = −12.6) (Table 1). A paired samples *t*-test showed significant difference (*t* = 3.384, df = 45, *P* = 0.001) between observed tile size selection scores and expected tile size selection scores with observed occurring more than expected. Linear regression showed no statistically significant relationship (*r*^2^ = 0.018, F_1,44_, *P* = 0.372) between scorpion length and tile size selection scores.

When treating all tiles as equally likely to be selected, *V. carolinianus* females from the 2014 France Mountain population also displayed a significant preference for larger tiles (*χ*^2^ = 30.831, *P* < 0.001, df = 2, expected *N* = 29.7). With tile surface areas included in analysis, no significant preference was found (*χ*^2^ = 0.691, *P* = 0.708, df = 2). Expected *N* values for small, medium, and large tiles were 12.3, 27.6, and 49.1, respectively. Residual values indicated that *V. carolinianus* selected small tiles slightly less than expected (residual = −1.3), medium tiles slightly less than expected (residual = −2.6), and large tiles slightly more than expected (residual = 3.9) (Table 1). A *t*-test revealed significant difference (*t* = 2.429, df = 29, *P* = 0.022) between observed tile size selection scores and expected tile size selection scores with larger tiles being selected more often than expected. Linear regression found no statistically significant relationship (*r*^2^ = 0.054, F_1,28_ = 1.611, *P* = 0.215) between scorpion length and observed tile size selection scores. A chi-square contingency table found no significant differences in surface area choice among the three populations of *Vaejovis carolinianus* (*χ*^2^ = 5.924, *P* = 0.2049, df = 4).

*Transect Data:* The majority of rocks in each transect fell into the Very Small (VS) category: Hwy 96 Roadcut = 87.99%, France Mountain firepit = 60.53%, and France Mountain logging trail = 70.97% (Figs. 2, 3, and 4). Due to relatively low numbers of occupied rocks (those being used as retreat sites), rock measurements from all three locations (four transects) were combined in order to compare sizes of unoccupied rocks to occupied rocks. Significant differences were found between the two groups (*n*_unoccupied_ = 730, *n*_occupied_ = 17, *U* = 554.5, *P* < 0.001) with occupied rocks being larger than unoccupied rocks, but no significant differences existed among rock sizes from the three locations (*χ*^2^ = 2.789, *P* = 0.248, df = 2, with mean surface area rank of 364.70 for Hwy 96 roadcut, 421.76 for France Mountain firepit, and 380.8 for France Mountain Logging Trail).

**Figure 2:**
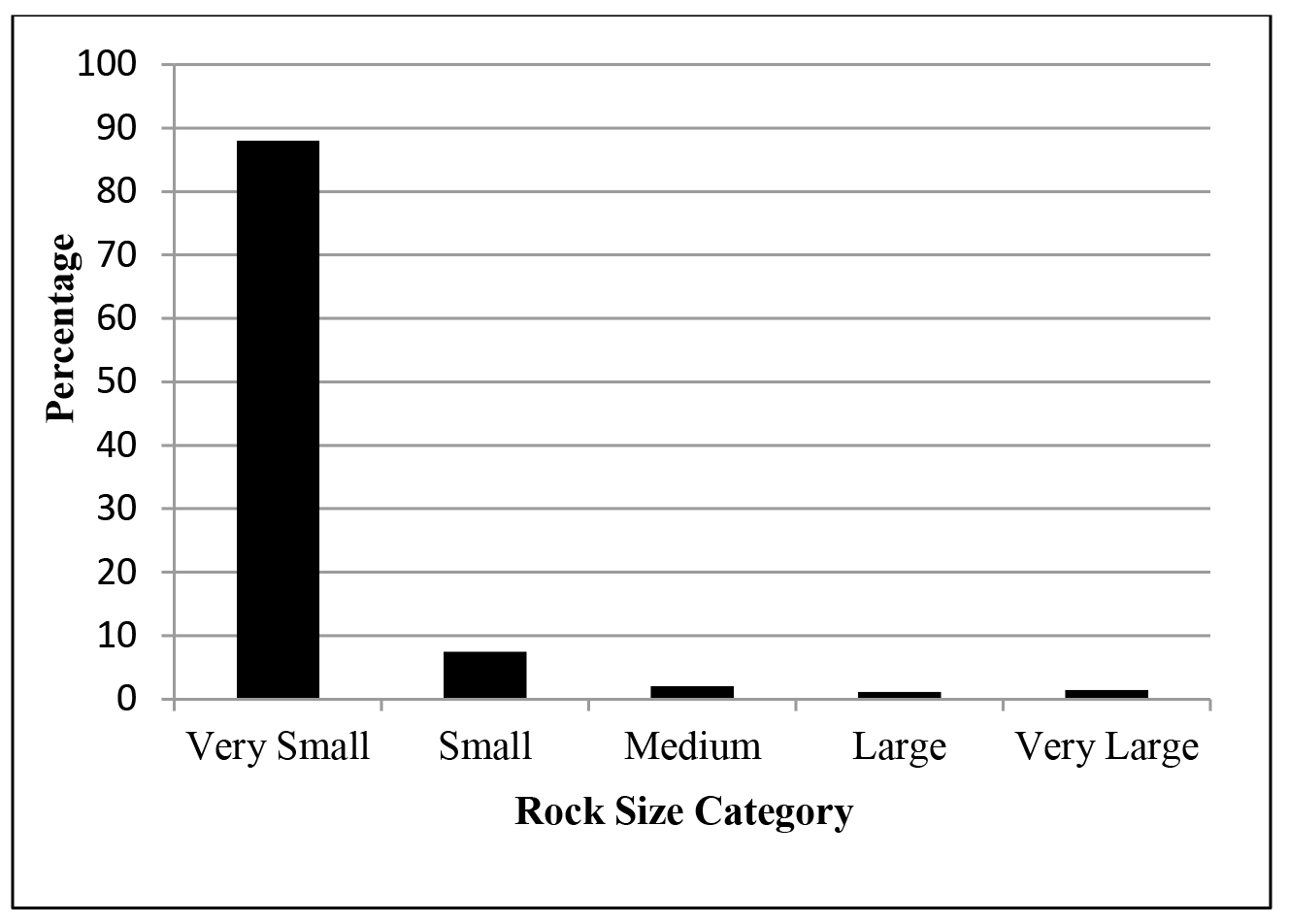
Breakdown of rock sizes within transect established at Hwy 96 roadcut. The graph shows that the majority of rocks were Very Small (87.99%) and considered too small to be selected for a retreat site.

**Figure 3:**
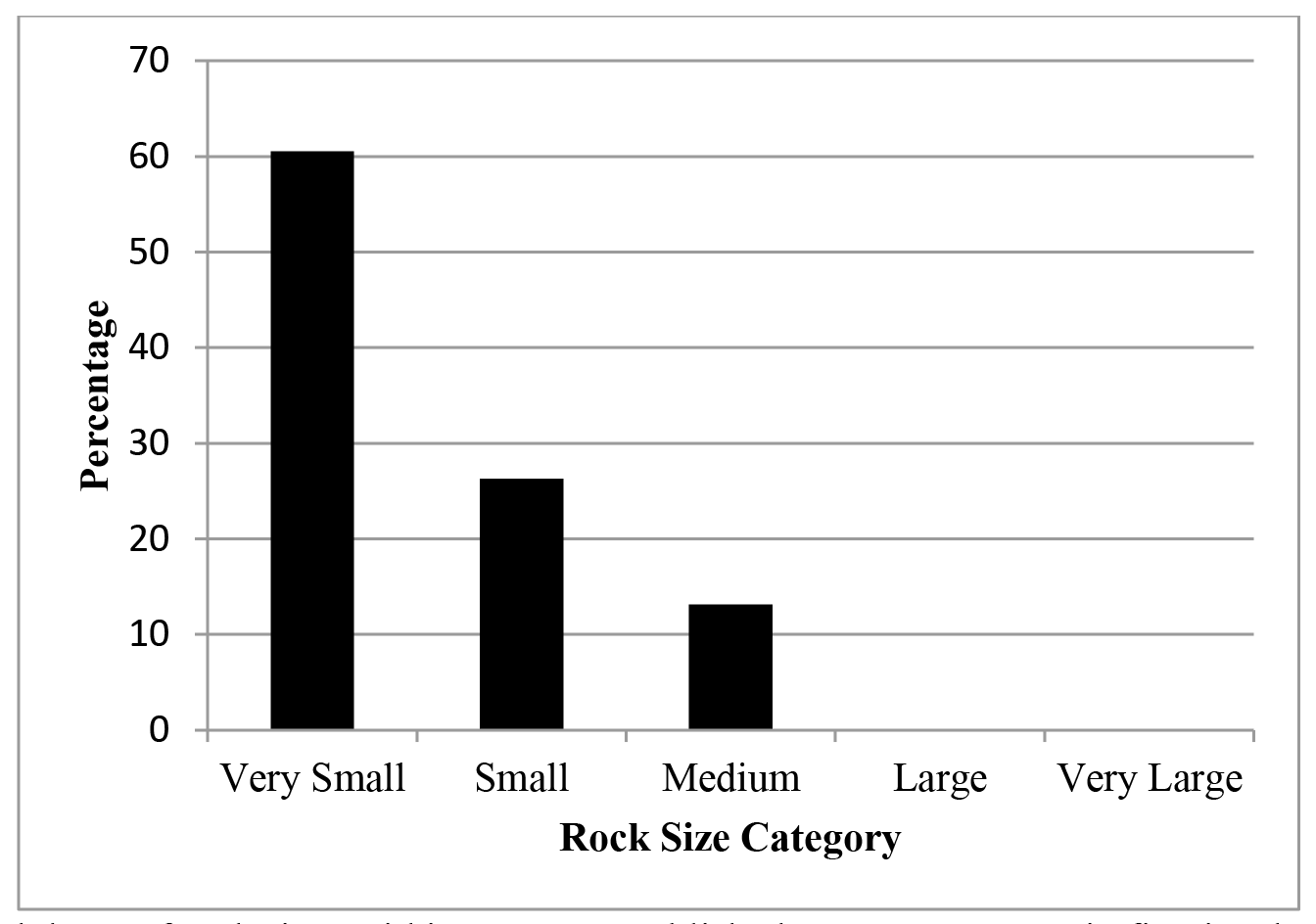
Breakdown of rock sizes within transect established at France Mountain firepit. The graph shows that the majority of rocks were Very Small (60.53%) and considered too small to be selected for a retreat site.

**Figure 4:**
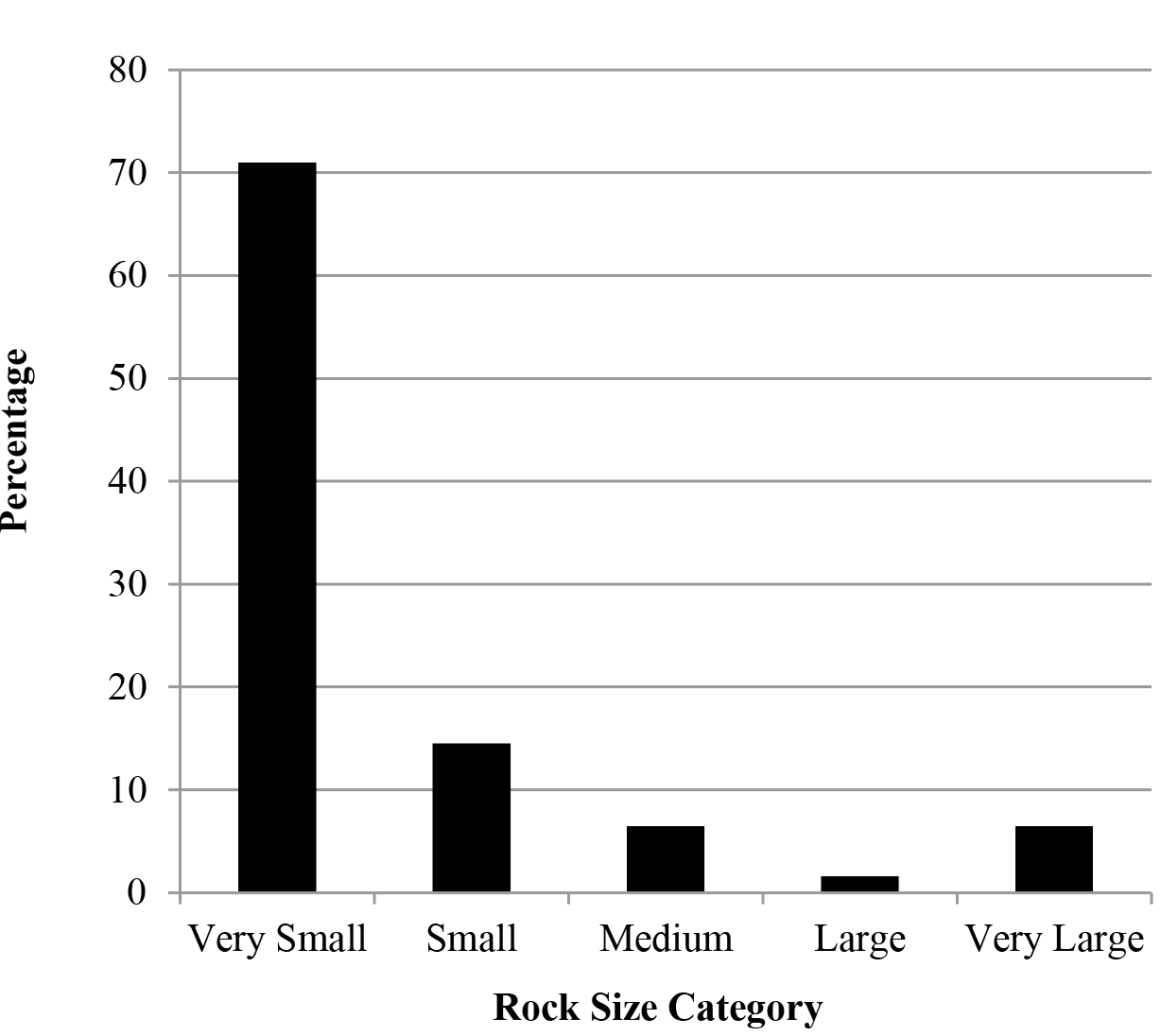
Breakdown of rock sizes within transect established at France Mountain logging trail. The graph shows that the majority of rocks were Very Small (70.97%) and considered too small to be selected for a retreat site.

Of 732 rocks turned (all four transects combined), only eight were occupied (three very large, one medium, four small). Nine additional scorpions were found while randomly selecting and flipping rocks outside the transect at the Hwy 96 roadcut site. Three occupied rocks were in the small size class, one in medium size class, and five were in the very large size class. I used a *G*-test to compare number of occupied rocks in each size class to total number of rocks available in each size class. I omitted the VS category (rocks too small to be used as retreat sites) and the large category (no large rocks were found to be occupied) from this analysis. *Vaejovis carolinianus* displayed significant preference among the three utilized sizes of cover objects (*χ*^2^ = 18.017, *P* < 0.001, df = 2). Expected *N*values were 11.2 for small rocks, 3.7 for medium rocks, and 2.2 for very large rocks. Very large rocks were utilized almost four times as much as expected (observed *N* = 8, residual = 5.8), and medium rocks were utilized approximately half as much as expected (observed *N* = 2, residual = −1.7). Small rocks were utilized less than half as much as expected (observed *N* = 7, residual = −4.2).

*Moisture Content of Soil at Retreat Sites:* Population sizes for these trials were as follows: Hwy 96 roadcut (*n* = 22), 2013 France Mountain (*n* = 40), and 2014 France Mountain (*n* = 30). I excluded two scorpions from the Hwy 96 roadcut population and five scorpions from the 2013 France Mountain population from analyses since they were found elsewhere in the tray on a single trial day. *Vaejovis carolinianus* from the Hwy 96 roadcut displayed no significant preferences among the three moisture levels (*χ*^2^ = 2.452, *P* = 0.294, df = 2) with an expected N of 20.7 (Table 2; Fig. 5). A *t*-test comparing observed soil moisture selection scores to expected soil moisture selection scores revealed no significance (*t* = 0.621, df = 21, *P* = 0.541). Linear regression comparing scorpion length to observed soil moisture selection scores showed no significant relationship (*r*^2^ = 0.107, F_1,21_: = 2.391, *P* = 0.138).

**Table 2:** Observed and expected *N* values with residual values (observed *N* - expected *N*) from chi-square tests on soil moisture selection.

| Population | Moisture Level | Observed $N$ | Expected $N$ | Residual |
| --- | --- | --- | --- | --- |
| Hwy 96 Roadcut | Dry | 20 | 20.7 | -0.7 |
|  | Damp | 16 | 20.7 | -4.7 |
|  | Wet | 26 | 20.7 | 5.3 |
| 2013 France Mtn | Dry | 44 | 38.3 | 5.7 |
|  | Damp | 40 | 38.3 | 1.7 |
|  | Wet | 31 | 38.3 | -7.3 |
| 2014 France Mtn | Dry | 19 | 30.0 | -11.0 |
|  | Damp | 31 | 30.0 | 1.0 |
|  | Wet | 40 | 30.0 | 10.0 |

For the 2013 France Mountain population, *V. carolinianus* displayed no significant preferences among the three moisture levels (*χ*^2^ = 2.313, *P* = 0.315, df = 2) with an expected *N* = 38.3 (Table 2). Significance was found (*t* = −2.047, df = 39, *P* = 0.047) from the *t*-test comparing observed soil moisture selection scores to expected soil moisture selection scores with dry soils selected more often than expected. Linear regression comparing scorpion length to observed soil moisture selection scores did not show any significant relationship (*r*^2^ < 0.001, F_1,38_ = 0.007, *P* = 0.934).

*Vaejovis carolinianus* from the 2014 France Mountain population displayed significant preference among the three moisture levels (*χ*^2^ = 7.400, *P* = 0.025, df = 2) with an expected *N* = 30 (Table 2). Wet soil was selected more than expected (residual = 10.0), damp soil was selected slightly more than expected (residual = 1.0), and dry soil was selected less than expected (residual = −11.0). Significance was found (*t* = 2.429, df = 29, *P* = 0.022) when comparing observed soil moisture selection scores to expected soil moisture selection scores with wet soils being selected more than expected. No significant relationship (*r*^2^ = 0.049, F_1,28_ = 1.431, *P* = 0.242) was found in a linear regression comparing observed soil moisture selection scores to scorpion length.

**Figure 5:**
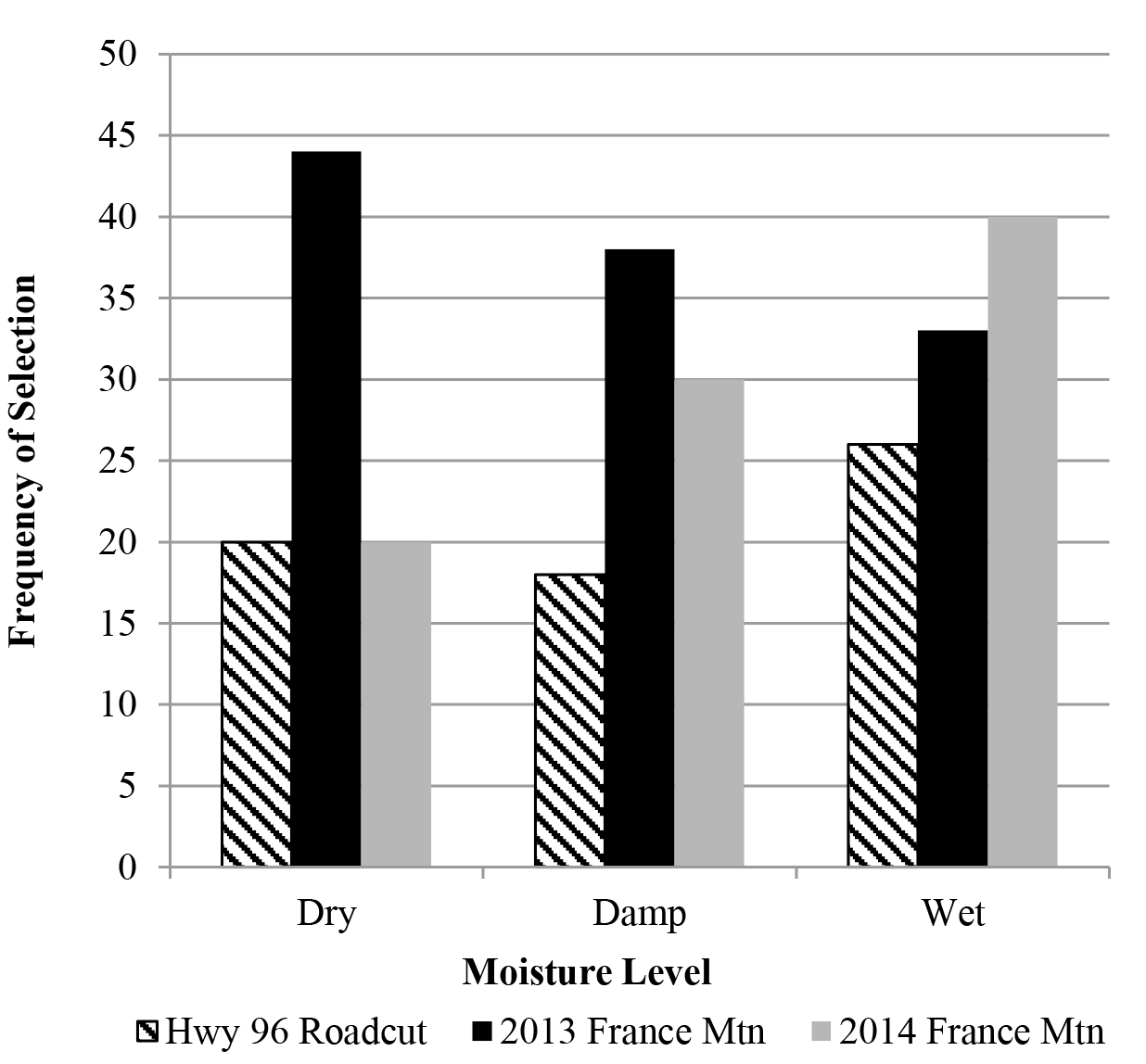
Frequency of selection of each of three soil moisture levels by three populations of the scorpion *Vaejovis carolinianus.*

## DISCUSSION

Animals usually face a number of important, yet conflicting demands during the course of a day (Downes and Shine, 1998). Predator avoidance and resource acquisition are of prime importance, but ectotherms must also seek retreat sites with conditions satisfactory to aid in thermoregulation. In this study, I examined two factors that may influence retreat site choice by the scorpion *Vaejovis carolinianus*: retreat site surface area and soil moisture.

Results of laboratory trials and transect measurements indicate *V. carolinianus* does not select retreat sites based on surface area. All three populations showed significant preference for large tiles when tile surface area was not considered during statistical analyses. When surface area was included in analyses, selection of the large tile did not occur more than expected. In other words, scorpions were more likely to be found under the larger tiles but did not appear to exhibit a clear preference for them. Lab results are somewhat consistent with field results in that both show that scorpions preferred larger size objects as retreat sites, but statistical significance is much stronger with field data. Although overall surface area of smaller rocks was greater (due to quantity), scorpions seemed to prefer larger rocks as retreat sites despite being far less common. Larger rocks provide greater protection from predators, sunlight, and desiccation, but the cost of this protection is decreased access to prey items.

Retreat site availability and general habitat differed somewhat between the two sites. Quantity of potential retreat sites was greater at the Hwy 96 roadcut, located between a bluff face and Hwy 96. The nearest wooded area to the capture location as at the top of the bluff. France Mountain capture sites were located either right at the forest edge (logging trail) or less than 15 m from the forest edge (firepit). The amount of low-lying vegetation near potential retreat sites was noticeably lower at France Mountain capture sites. This may affect retreat size choice, since *V. carolinianus* commonly resides near the edge of moist forest habitats (Benton, 1973).

Due to relatively low daily temperature fluctuation (~10°C) in the Upper Cumberland region, rock thickness was not taken into consideration during field and lab trials on surface area size. Low temperature fluctuation may explain why *V. carolinianus* were found under small and medium-size rocks as well as large rocks. A study in the western U.S. included surface area and thickness in comparisons involving thermal properties of retreat sites selected by *Vaejovis electrum* and *V. cashi* (Becker, 2013). Both species chose rocks larger and thicker than the mean for the habitat, possibly because larger, deeper rocks have more stable temperatures underneath. These *Vaejovis* species are found in areas that have a wide range in daily temperature fluctuation, so thermal properties of retreat sites (including rock thickness) would be of great importance since scorpions are ectothermic and thermoregulate by changing their behavior or location.

Almost all scorpions captured for my study were found under rocks resting directly on moist soil, so it was anticipated that soil moisture was an influential factor in retreat site selection. However, lab results did not agree with field observations since no significant preference was found for soil moisture content. Results for the Hwy 96 roadcut population and the 2013 France Mountain population showed no significant soil moisture preference. The 2014 France Mountain population showed statistical significance, indicating a preference for wetter soils. Time in captivity was probably the main reason for the difference, because the 2013 France Mountain population had been supplied with fresh water regularly while the 2014 France Mountain population, being newly caught, was still accustomed to dry conditions of their normal habitat. Benton (1973) conducted a study examining humidity preference with *V. carolinianus.* He learned that humidity preferences change based on time of day and they seem to follow the same “clock” that regulates circadian rhythm. Benton (1973) concluded that *V. carolinianus* preferred 98 – 100% relative humidity diurnally, and can distinguish between 64% and 98% humidity. No relative humidity preference was demonstrated nocturnally. Substrate moisture is a potential water source for scorpions, while others include fluids of captured prey, bulk surface water, atmospheric moisture, and metabolic water (Hadley, 1974). Scorpions lose water at a relatively low rate (Gefen and Ar, 2004) and do not leave the retreat site every night, so it would appear that substrate moisture and metabolic water would be two important water sources while scorpions are at retreat sites. Another important reason to select retreat sites based on soil moisture levels would be ability to minimize water loss.

Suggestions for future studies involving *V. carolinianus* are numerous, for little is known about this seldom-researched species. More research needs to be conducted with retreat site selection; for example, studies could be conducted regarding thermal properties of retreat sites. Other suggested studies include additional research on male *V. carolinianus,* as well as juveniles. Currently, all that seems to be known about males is (1) they do not aggregate, and (2) they prefer a different type of microhabitat as a retreat site than do females. Future studies should also examine the maternal costs of parturition and rearing of young, as well as food preferences and growth rate of juveniles.

## Acknowledgements

I thank my advisor Dr. Christopher A. Brown for his advice and encouragement. I also thank Margaret “Maggie” Melendez for her assistance in scorpion care, and I thank my parents for their support and encouragement throughout my academic career.

